# Invasion and migration of spatially self-limiting gene drives: a comparative analysis

**DOI:** 10.1101/159855

**Authors:** Sumit Dhole, Michael R. Vella, Alun L. Loyd, Fred Gould

## Abstract

Recent advances in research on gene drives have produced genetic constructs that could theoretically spread a desired gene (payload) into all populations of a species, with a single release in one place. This attribute has advantages, but also comes with risks and ethical concerns. There has been a call for research on gene drive systems that are spatially and/or temporally self-limiting. Here we use a population genetics model to compare the expected characteristics of three spatially self-limiting gene drive systems: one-locus underdominance, two-locus underdominance, and daisy-chain drives. We find large differences between these gene drives in the minimum release size required for successfully driving a payload into a population. The daisy-chain system is the most efficient, requiring the smallest release, followed by the two-locus underdominance system, and then the one-locus underdominance system. However, when the target population exchanges migrants with a non-target population, the gene drives requiring smaller releases suffer from higher risks of unintended spread. For payloads that incur relatively low fitness costs (up to 30%), a simple daisy-chain drive is practically incapable of remaining localized, even with migration rates as low as 0.5% per generation. The two-locus underdominance system can achieve localized spread under a broader range of migration rates and of payload fitness costs, while the one-locus underdominance system largely remains localized. We also find differences in the extent of population alteration and in the permanence of the alteration achieved by the three gene drives. The two-locus underdominance system does not always spread the payload to fixation, even after successful drive, while the daisy-chain system can, for a small set of parameter values, achieve a temporally-limited spread of the payload. These differences could affect the suitability of each gene drive for specific applications. **Note:** This manuscript has been accepted for publication in the journal Evolutionary Applications and is pending publication. We suggest that any reference to or quotation of this article should be made with this recognition.

## Introduction

Gene drives are genetic constructs that can be used to spread a desired gene, often referred to as the “payload”, into a sexually reproducing population even when the inserted construct or the payload reduces the fitness of the individual carrying it (Curtis 1968; Burt 2003; reviewed in Sinkins and Gould 2006; Alphey 2014). This ability to spread genes that reduce individual fitness has potential for use in controlling disease vectors and agricultural pests as well as in conservation biology (Gould 2008; Esvelt et al. 2014). For instance, a gene drive that spreads a payload gene that drastically reduces female fecundity could be used to control mosquito populations (Alphey et al. 2002; Burt 2003; Deredec et al. 2008). Other possibilities include genetic constructs that cause a strong skew in the offspring sex ratio toward males (Galizi et al. 2014; Galizi et al. 2016) or that interfere with a vector's ability to transmit a pathogen (Gantz et al. 2015). Even in cases where the goal of introducing a new gene is not aimed at reducing fitness, there are often some fitness costs associated with the introduction of a novel genetic construct (Franz et al. 2014).

With recent advances in genome editing utilizing CRISPR-Cas9, the efficiency of constructing gene drives has substantially improved, and experiments have recently been performed that demonstrate successful gene drive in laboratory populations of mosquitoes and Drosophila with the new constructs (e.g. Galizi et al. 2016; Hammond et al. 2016; Gantz and Bier 2015; Gantz et al. 2015; Champer et al. 2017a; Champer et al. 2017b). These gene drives have multiple advantages over certain traditional methods of population alteration. Because gene drives only spread through sexual reproduction between individuals of a population, their effects are expected to be species-specific, unlike the impacts of synthetic insecticides that are commonly used today and affect a broad array of species. Additionally, gene drives are expected to perpetuate themselves, so follow-up costs after release of an engineered strain could be very low.

However, like other powerful technologies, gene drives are not free of potential risks. With the ability of gene drives to spread among populations connected by even limited gene flow, a thorough understanding is needed of the possible risks involved with spread beyond the intended population, either through natural migration of transgenic organisms from the targeted population or through humans intentionally or unintentionally transporting transgenic individuals long distances (Marshall and Hay 2012; NASEM 2016). Indeed, some gene drives, such as homing drives (HDs) based on genetic constructs with CRISPR-Cas9, may be capable of indefinite spread across the whole geographic range of a species and that of any other species with which they are able to mate and produce some offspring (Burt 2003; Deredec 2008; Alphey and Bonsall 2014). Uncontrolled spread of a gene drive may pose environmental risks depending upon the effect of the gene drive and the payload on the species. Indefinite spread could also violate the Cartagena Protocol (CBD 2016) if the spread occurs across international political boundaries without consent (NASEM 2016). Gene drives that can achieve localized population alteration are therefore highly desirable in many contexts. Easy reversibility of a gene drive would add a further safety net, if the population needs to be restored to a wild-type state after release of the gene drive.

Certain proposed types of gene drive are theoretically expected to spread in a population only if they are released above a certain threshold frequency relative to the naturally occurring individuals in a population (Curtis and Robinson 1971; Davis et al. 2001; Magori and Gould 2006; Buchman et al. 2016). Gene drives with a high threshold introduction frequency will require a larger introduction of transgenic individuals for successful population alteration, and are likely to be economically less efficient. However, if the threshold introduction frequency for a gene drive is higher than the frequency that could be attained by migration or accidental release, such gene drives may provide an approach for localized population alteration and will raise fewer concerns about disruption of ecological systems.

Of the gene drives proposed for localized population alteration (Gould et al. 2008; Rasgon 2009; Marshall and Hay 2014; Buchman et al. 2016), one‐ and two-locus engineered underdominance seem theoretically promising, and have been built and shown to be capable of drive in engineered laboratory populations (Akbari et al. 2013; Reeves et al. 2014). Recently, Noble et al. (2016) proposed an intriguing new structure for a localized, temporary gene drive, and it has gathered much attention from the scientific community (Camporesi and Cavaliere 2016; Harvey-Samuel et al. 2017; Unckless et al. 2017), as well as from popular media (New Scientist, June 2016; The Verge, June 2016; Epigenie, August 2016). This new gene drive, called the 'daisy-chain driveâ€™, is a form of a homing drive (HD) based on CRISPR-Cas 9 (Noble et al. 2016). Unlike previously proposed HDs, the daisy-chain system separates the elements of a simple HD onto independently segregating loci (Noble et al. 2016). This separation of the elements was theoretically predicted to result in a drive that is localized, and would restrict the population alteration to a limited time period (Noble et al. 2016).

Although these aforementioned gene drives have theoretically been shown to be capable of localized spread under certain conditions, it is not clear how they compare to each other under different conditions, especially in their level of localization and permanence. Here we present a comparative analysis of the level of localization of three gene drives – the daisy-chain drive, and the two-locus and one-locus engineered underdominance drives. We address the efficiency of these gene drives to spread a payload allele in an isolated population, as well as their ability to bring about local population alteration in face of unidirectional incoming and bidirectional migration with a neighboring population. Our goal here it to compare the behavior of these gene drives under a broad set of identical conditions.

We use a population genetic model to simulate the spread of a payload allele, which is linked to one of the components of the gene drive, and which may incur a fitness cost on the individual bearing it. We focus mainly on systems where the drive is aimed at altering the characteristics of a population (e.g. inability to transmit a pathogen) and have relatively low levels of fitness cost. However, we also adapt our model for an initial comparison of these gene drives when the goal is to suppress a population. We find large differences in the efficiencies as well as the levels of localization of the three gene drives. We discuss the relative suitability of the different gene drives for applications in different scenarios.

## Model

We use population genetic models to separately simulate the spread of a desired “payload” allele with each of the three types of gene drives in an isolated diploid population. We also compare the within and among population spread of the payload allele by the three gene drives when the target population exchanges migrants with a neighboring population.

We follow the spread of the payload allele after a single initial introduction of individuals homozygous for the complete drive constructs into the target population. The payload allele fitness cost can be varied depending upon whether the gene drive is intended for population suppression or for population replacement. The fitness cost associated with being homozygous for the payload allele is given by *s_p_*, while the cost to heterozygotes is modified for specific dominance patterns of the payload allele. We show results in the main text only for multiplicative cost of the payload allele. Results with additive, recessive and dominant payload costs are provided in the online Appendix 1.

We model a lifecycle with non-overlapping generations. We assume that the population is very large, and model only gene frequency-based dynamics (no multi-generation density-dependent dynamics). The gene drives addressed here differ in the mechanisms that result in their spread in a population – the daisy-chain drive relies on homology-directed repair (HDR) following double-stranded breaks in the DNA, while the underdominance drives use lower fitness (or lethality) of transgene-wildtype heterozygotes for their propagation. The variation in genotypic fitnesses caused by these driving mechanisms results in natural selection. The payload allele may further add to variation in genotypic fitnesses. In our model, natural selection due to the drive mechanism and due to the fitness costs imposed by the payload allele occurs in immature stages or in adults before reproduction. Migration occurs after natural selection and is followed by reproduction within a population. Individuals in our model mate randomly with respect to the alleles at the gene drive loci. Offspring are produced following free recombination.

### Gene drive structures

Below we describe the structure of the three gene drives that we compare. The constructs used to build each of the gene drives can be modified to exhibit a large variety of properties, such as tissue-specific or sex-specific patterns of expression, condition-dependent fitness costs, etc. For the comparative analysis here, we mainly model the basic forms of these gene drives. But, for the comparison of the capacity for population suppression, we model these drives with female-limited payload gene expression. We also assume that the payload allele is carried at only one locus and has the same phenotypic expression with each gene drive.

#### Daisy-chain drive

The daisy-chain drive is a type of CRISPR-based homing drive (Esvelt et al. 2014). We model a 3-element daisy-chain drive as described by Noble et al. (2016). In this daisy-chain drive each driving element and the element that it drives are located at independently segregating loci (Noble et al. 2016; Figure 1). In a 3-element daisy-chain drive, a payload allele at locus A is driven by a second, independently segregating allele at locus B, which in turn is driven by a third independently segregating allele at locus C (see Figure 1). The third element is not driven and is expected to decline quickly. The separation of the drive elements is intended to prevent an indefinite spread of the gene drive. The loci A and B are chosen to be carrying genes essential for survival (or reproduction), and are haplo-insufficient. This design, in combination with the multiple double-stranded cuts, is expected to reduce the likelihood of appearance of genotypes that are resistant to the drive mechanism (Esvelt et al. 2014; Noble et al. 2016; Noble et al. 2017), but has not yet been empirically demonstrated (but see Champer et al. 2017a). The transgenic allele at the B locus is a modified version of the corresponding essential wild-type gene, while the transgenic allele at the A locus could be either a modified functional or dysfunctional variant based on whether the goal is population replacement or suppression.

**Figure 1:**
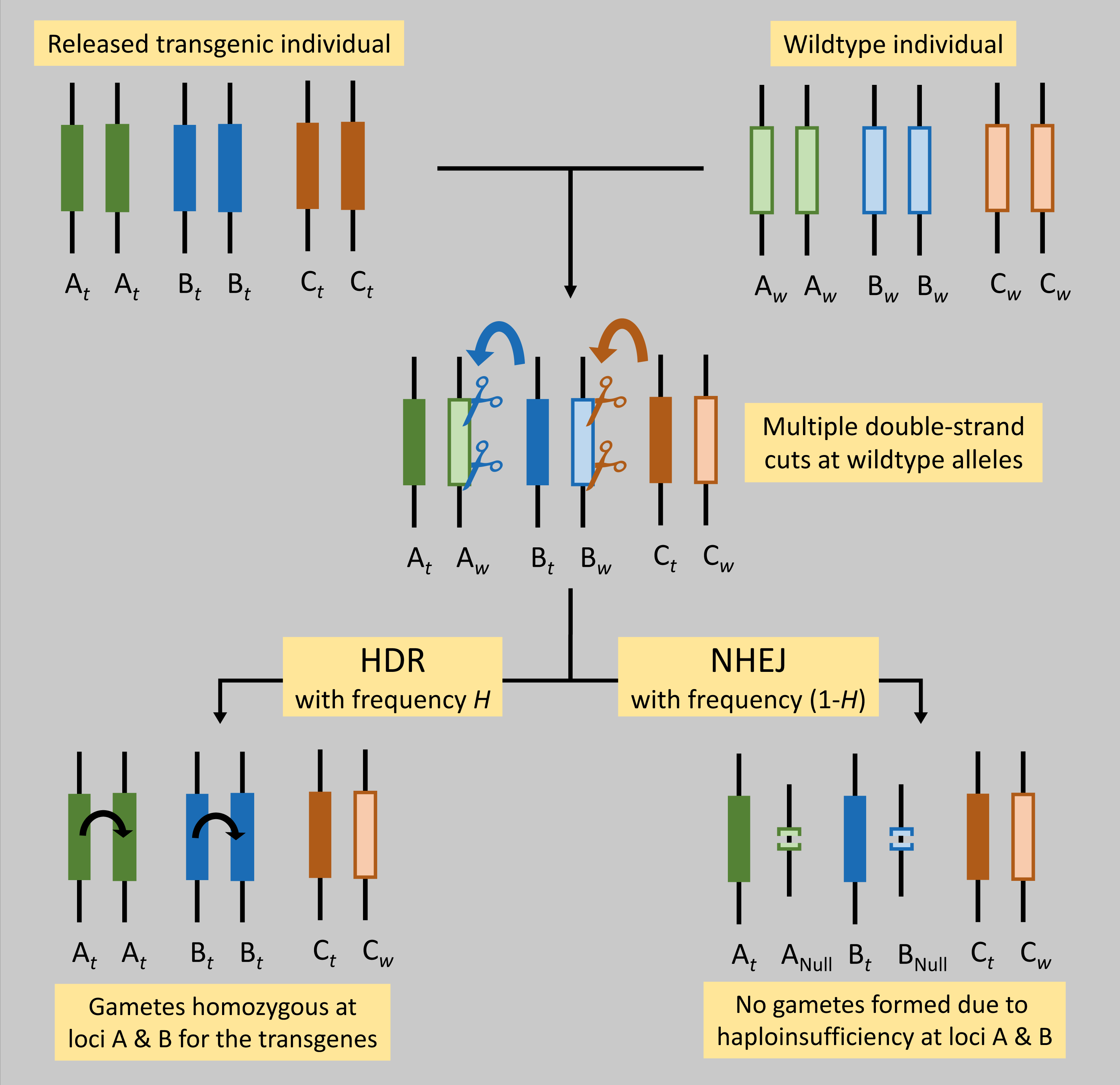
The Daisy-chain drive structure (adapted from Noble et al. 2016) and its mechanism are illustrated. Parental genotypes are shown in the top row. Transgenic alleles are shown with a subscript ‘*t*’ and with darker colors, while wild-type alleles have subscript w and lighter colors. The payload gene is located within the A*_t_* allele. The transgenic alleles are expressed in the germline cells of the offspring (middle row) resulting in multiple double-stranded breaks on the wild-type alleles. Repair through non-homologous end-joining (NHEJ) results in deletions on the wild-type chromosomes and failure of gametogenesis (or production of gametes that fail to produce viable offspring) due to haploinsufficiency of the genes, whereas homology directed repair (HDR) restores a full complement of the genome and enables successful gametogenesis.

We use a slightly modified version of the notation used by Noble et al. (2016), because we use a discrete-time model (unlike the continuous time version they use) and also because our notation allows easier comparison with the notation for the other gene drives presented here. The dynamics of the gene drive are not altered by the differences in notation.

Following Noble et al. (2016), there are two possible functional alleles at each of the three loci – a transgenic allele (represented by subscript ‘*t*’) and a wild-type allele (represented by subscript ‘*w*’). The B*_t_* allele is dominant and drives the A*_t_* allele by making multiple double-stranded cuts at all copies of the wild-type A*_w_* allele (but leaving the A*_t_* allele intact). In heterozygous cells, the cut at the A*_w_* allele can be repaired through homologous recombination (homology-directed repair or HDR) using the uncut A*_t_* allele as a template, thus rendering the cell homozygous for the A*_t_* allele after repair. Similarly, the C*_t_* allele is dominant and converts B*_t_*B*_w_* heterozygotes to B*_t_*B*_t_* homozygotes. HDR can occur only in heterozygotes. Homozygotes for the wild type allele at locus A and B can only be repaired through non-homologous end joining (NHEJ) that would result in a deletion at the location of the multiple cuts, resulting in non-functional alleles.

In the 3-element daisy-chain drive, the A*_t_* allele includes a payload gene sequence. This payload can alter an individual's characteristics (e.g. not vectoring a pathogen) or decrease fitness. The drive components of the transgenic alleles at the B and C loci are, by design, only expressed in germline cells, so that homing only occurs in the germline. It is assumed that even a single copy of a B*_t_* allele always results in complete cutting of all A*_w_* alleles in the cell (i.e., the B*_t_* allele is completely dominant). Similarly, a C*_t_* allele always cuts all copies of the B*_w_* allele. The frequency of successful HDR after a wildtype allele is cut is referred to as ‘homing efficiencyâ€™, and is denoted by the parameter H. In absence of HDR after a cut, NHEJ results in a non-functional allele and the germline cell fails to produce gametes (or produce gametes that fail to produce viable offspring) due to haploinsufficiency of the genes. An individual with genotype A*_t_*A*_w_* and at least one copy of the B*_t_* allele would thus lose a fraction (1-H) of its functional gametes, and only produce H functional gametes relative to an individual that does not have any cuts occurring. B*_t_*B*_w_* individuals pay a similar cost when they also inherit a copy of the C*_t_* allele. As in Noble et al. (2016), the cost of such driving at the two loci is assumed to be multiplicative; i.e. A*_t_*A*_w_*B*_t_*B*_w_*C*_t_*C*_w_* individuals produce H^2^ functional gametes relative to completely wild-type individuals. This is the cost of the drive mechanism, and is independent of the cost of the phenotypic effect of the payload allele (A*_t_*).

The CRISPR-based gene drives have been shown to exhibit homing efficiencies as high as 99% (Gantz et al. 2015; Hammond et al. 2016). For the purpose of the analysis here we use a homing efficiency (H) of 95% throughout, unless noted otherwise.

To describe the genotypic relative fitnesses, we denote the diploid genotypes at the A locus A*_w_*A*_w_*, A*_w_*A*_t_* and A*_t_*A*_t_* simply as A0, A1 and A2, respectively. Genotypes at the B and C loci are denoted in a similar fashion. Thus the complete diploid genotypes can be written as A*_x_*B*_y_*C*_z_*, where x, y and z can each have values 0, 1 or 2 for genotypes ww, tw and tt, respectively. We then define the following function as the component of fitness attributed to the driving action of the alleles at the B and C loci.

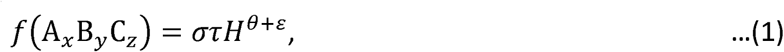

where σ,τ,θ and *ε* are modifiers that alter the function to give fitnesses of the different genotypes depending upon whether cutting (and then HDR) occurs at each locus. The terms σ and τ modify the equation to account for whether cutting occurs at a loci B and A, respectively, and also whether a transgenic template is available for HDR when cutting does occur. The modifiers θ and *ε* modify the cost of inefficient homing for zero, one or two loci. Their values for the different genotypes are

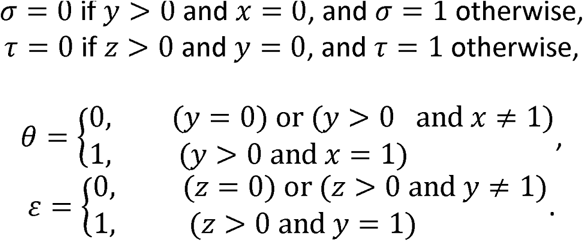

As mentioned previously, the payload allele confers a homozygous fitness cost s*_p_*. Here we describe genotypic fitness assuming multiplicative cost of the payload. Equations for other patterns of payload costs are given in Appendix 1. The component of fitness of genotype A*_x_*B*_y_*C*_z_*, that is attributed to the payload allele is given by,

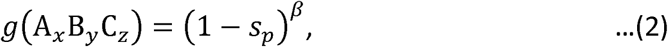

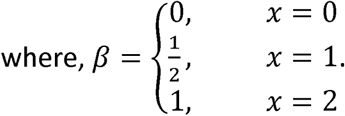

The relative fitness of a genotype A*_x_*B*_y_*C*_z_* after selection due to the drive mechanism and due to the payload allele is then given by the product,

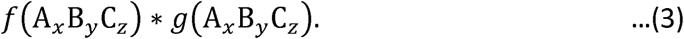

In the analyses shown here we assume that the non-payload transgenic alleles do not impose any direct fitness costs, besides the costs that arise through the drive mechanism. Analyses with additional fitness costs of the non-payload elements are provided in Appendix 2.

#### Underdominance

Davis et al. (2001) described two engineered underdominance gene drives, each with two engineered constructs, located either at a single locus or at two separate loci. The two constructs are alleles at a single locus in the one-locus underdominance system, while in the two-locus underdominance system, the two constructs are alleles at two independently segregating loci.

The **One-locus underdominance** drive is modeled to consist of two engineered constructs, each of which is composed of four tightly linked elements – 1) a gene coding for a lethal toxin, 2) a cis-promoter for the toxin gene, 3) a suppressor specific for the other construct's promoter and 4) a payload gene (Figure 2A).

**Figure 2:**
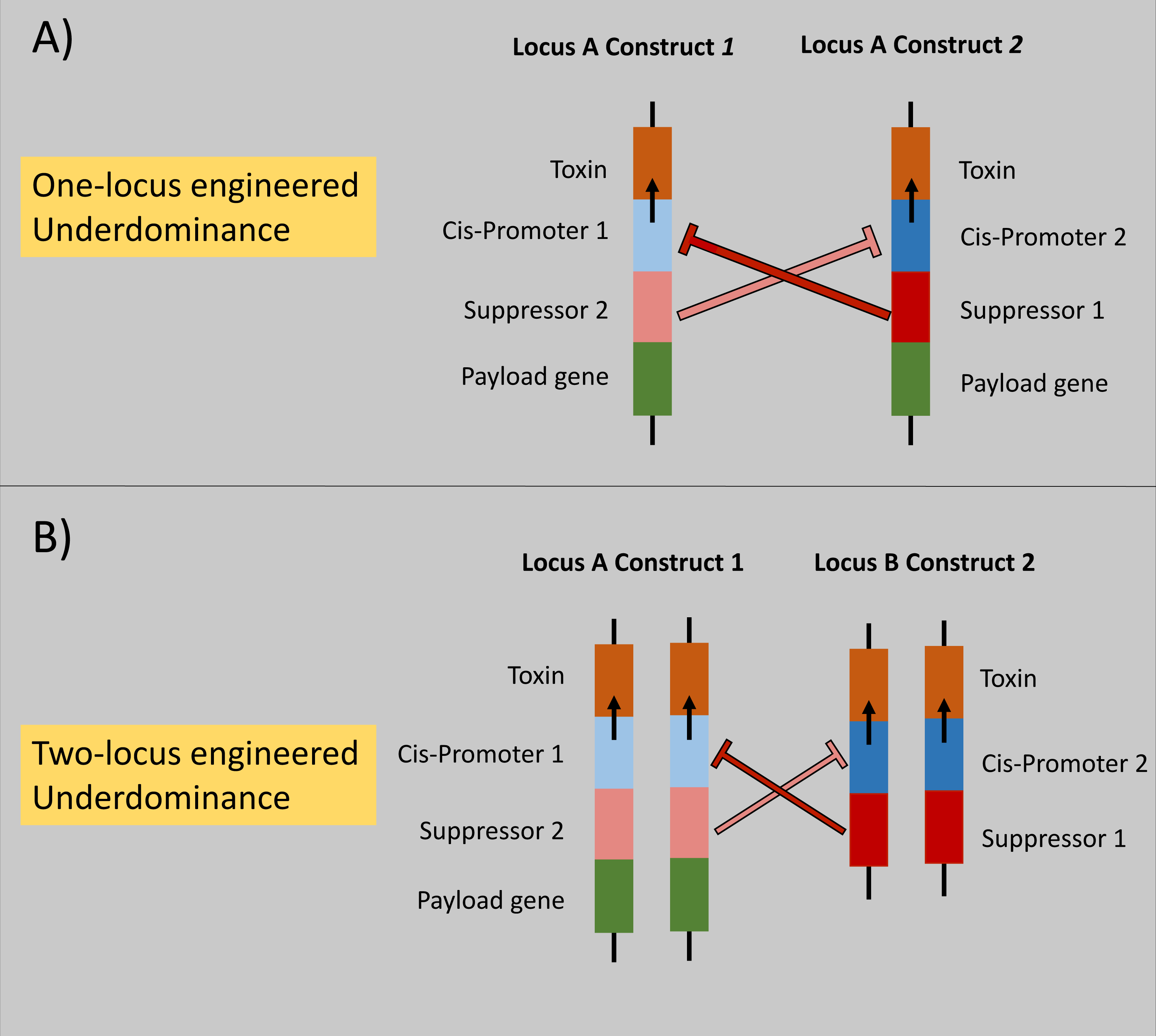
The structure of the two types of underdominance drives is illustrated. Only transgenic individuals that would be released are shown. Suppressors on each construct block the promoters on the other construct.

The elements within each construct are tightly linked by design, and therefore each construct can be modeled as an allele at a single locus ‘A’. The two constructs are referred to as alleles A*_1_* and A*_2_*. Wild-type alleles at the locus A are denoted as A*_0_*. Individuals that inherit one copy of each transgenic allele (genotype A*_1_*A*_2_*) are viable, because the suppressor on each construct blocks the cis-promoter on the other allele, thus preventing the production of the toxin (see Figure 2A). Individuals that carry only one of the constructs (genotypes A*_1_*A*_1_*, A*_0_*A*_1_*, A*_2_*A*_2_* and A*_0_*A*_2_*) are not viable, as the toxin is not suppressed. Wild-type homozygotes (genotype A*_0_*A*_0_*) are, of course, viable. The expression of the toxins on each construct can be timed to occur at any desired life stage. The payload allele confers a multiplicative fitness cost of s*_p_* when homozygous. Note that for the one-locus underdominance drive the multiplicative cost pattern is indistinguishable from other dominance patterns (such as additive or dominant costs) because heterozygotes are not viable. The relative fitnesses of the different diploid genotypes for the one-locus underdominance system are given in Table 1.

**Table 1:** Relative fitnesses of diploid genotypes for the one-locus underdominance system.

| | $A_0$ | $A_1$ | $A_2$ |
| --- | --- | --- | --- |
| $A_0$ | 1 | 0 | 0 |
| $A_1$ | 0 | 0 | $(1 - s_p)$ |
| $A_2$ | 0 | $(1 - s_p)$ | 0 |

Table Note: Columns and rows show alleles inherited from each parent.

The **two-locus underdominance** drive is composed of constructs similar to those in a one-locus underdominance drive, but the two types of constructs are located at independently segregating loci (Figure 2B). The first construct is located at locus ‘A’, and locus ‘B’ carries the second construct. Each of the loci can thus have either a transgenic allele (denoted by subscript *t*) or a wild-type allele (denoted by the subscript *w*), with the allele A*_t_* representing construct 1 and allele B*_t_* representing construct 2. It is assumed that even one copy of a suppressor (heterozygous state at one of the loci) is sufficient to block all copies of the corresponding promoter on the other construct (see “strong [toxin] suppression” case in Edgington and Alphey 2017). Thus, only individuals with at least one copy of both transgenic alleles or no transgenic allele at all can survive. The payload allele can be linked to one or both of the constructs. For equivalence with the other two gene drives, we model a two-locus underdominance drive with the payload only linked to locus A (see Figure 2B). Note that the evolutionary dynamics of the two loci in such a two-locus underdominance model are symmetrical (Magori and Gould 2006) when the toxins on the two constructs result in identical fitness costs, therefore the choice of which of the two loci the payload allele is linked to does not change the results. The genotypic fitnesses for the two-locus underdominance system are shown in Table 2.

**Table 2:** Relative fitnesses of diploid genotypes for the two-locus underdominance system.

| | $A_w B_w$ | $A_w B_t$ | $A_t B_w$ | $A_t B_t$ |
| --- | --- | --- | --- | --- |
| $A_w B_w$ | 1 | 0 | 0 | $(1 - s_p)^{1/2}$ |
| $A_w B_t$ | 0 | 0 | $(1 - s_p)^{1/2}$ | $(1 - s_p)^{1/2}$ |
| $A_t B_w$ | 0 | $(1 - s_p)^{1/2}$ | 0 | $(1 - s_p)$ |
| $A_t B_t$ | $(1 - s_p)^{1/2}$ | $(1 - s_p)^{1/2}$ | $(1 - s_p)$ | $(1 - s_p)$ |

Table Note: Columns and rows show haplotypes inherited from each parent. Fitness is shown with multiplicative cost ( $s_p$ ) of the payload allele.

Note that although we describe the underdominance drives as containing a lethal toxin here, our model is equally applicable for describing the dynamics of underdominance drives that contain fertility-disrupting “toxins” that result in complete infertility instead of lethality. Fertility-disruption will however allow heterozygotes to reach adult stages, which may pose a problem for certain forms payload genes intended for disease-control.

### Migration

We assume adult individuals migrate after natural selection has occurred within each population, and that mating occurs immediately after migration. Moreover, we assume that migration is not influenced by individual genotypes at the gene drive loci. Thus, the group of emigrant individuals has the same genotypic frequencies as those in the source population after natural selection. We briefly discuss the potential effects of relaxing this assumption of genotype-independent migration in the Discussion section, but leave formal analysis for future studies.

#### Unidirectional migration to an island from the mainland

We first address a scenario of a targeted population on an island that receives migrants from the mainland each generation. The mainland population is assumed to be composed entirely of wild-type individuals, and so much larger than and far enough from the target population that no migration occurs from the island to the mainland (thus the mainland is assumed to always remain free of any gene drives). The island is modeled to receive an influx of wild-type emigrants at a rate given by the parameter *μ* from the mainland just before mating occurs every generation.

#### Bidirectional migration between two populations of similar size

In reality, migration is rarely completely unidirectional. To compare the level of localization of the different gene drives, we also analyze a scenario with bidirectional migration between the target population and a neighboring population of similar size.

We assume that the base migration rate ‘*μ*’ is equal in both directions, i.e. the same fraction of individuals migrate out of each population. When both populations have an equal number of fertile adults before migration, a fraction *μ* of each population after migration in each generation is composed of migrants from the other population. This population of emigrants and residents would form the breeding adult population in each generation.

Although we assume that the two populations initially have equal sizes, the fitness costs imposed by the gene drive can cause genetic load on the population, and reduce the fertile adult population size within a generation. If the two populations differ in size, this would lead to an asymmetry in the effective migration rate, or the immigration rate, because more individuals would leave the larger population than the smaller population (e.g. see Huang et al. 2011; North et al. 2013). Moreover, the individuals arriving from the larger population would make up a larger fraction of the smaller population. Thus the effective immigration rate would be asymmetric if the population sizes differ, even if the base migration rate is equal. To account for this, we use the genetic load generated in each generation to approximate the asymmetry in the effective immigration rates. The effective immigration rates in each direction in each generation are given by *μ_T_* for migration into the target population and *μ_N_* for migration into the neighboring population. These effective immigration rates are functions of the base migration rate and the relative genetic load on the two populations in a given generation. The genetic load represents a reduction in the population size from birth to fertile adults. The ratio of the sizes of the two fertile adult populations in a given generation then is given by

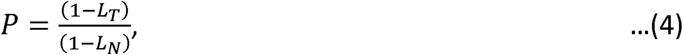

where L*_T_* and L*_N_* are the genetic loads in the target and the neighboring populations, respectively. The effective immigration rates” ar_e t“h en given by

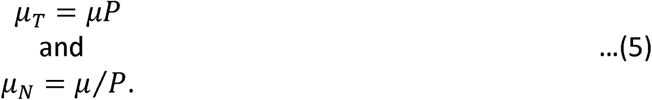

As may have been apparent above, for the purpose of the migration analysis, we assume that a reduction in the number of adults does not result in a reduction in the population size *across generations* (but, see the sections on population suppression in isolated populations below). This assumption is justified when density-dependent compensation occurs through changes in offspring production or density-dependent offspring survival. We recognize that such compensation may not be effective when the genetic load is extremely high. A more accurate estimation of the changes in the sizes of the two populations across generations would require explicitly incorporating density-dependent population dynamics, which would depend upon the species-specific ecological details (e.g. Alphey and Bonsall 2014). Given the scope of this study, and the fact that we find relatively low genetic loads across most of the conditions used for the migration analyses (see results on Suppression in isolated populations below), we leave long-term density-dependent dynamics for future studies.

We also perform this localization analysis without accounting for differences in population sizes, where the migration rate in either direction remains unchanged (remains equal to *μ*) irrespective of the genetic load on each population. This does not qualitatively change the results described below (see Appendix 3).

### Analyses

We simulate the spread of the gene drives after a single initial introduction of transgenic individuals into the target population. For the one-locus underdominance drive, transgenic heterozygotes (genotype X*_1_*X*_2_*) are introduced (see description above). For the two-locus underdominance and the daisy-chain drive, individuals homozygous for the gene drive are introduced. Our goal is to compare the efficiency of the three gene drives in spreading the payload allele into the target population, the level of localization of the payload allele to the target population (when bidirectional migration occurs between the target and the neighboring populations), and the degree of permanence of the payload gene in the targeted population. To this end, we calculate the mean frequencies (calculated each generation at birth) of the payload allele in both the target and the neighboring populations in the first 100 generations after the initial introduction (the payload allele approaches equilibrium frequencies within 50 generations in most cases). These mean frequencies are used for comparing the level of population alteration caused by the different gene drives in the target and the neighboring populations. The mean payload allele frequency gives a better measure of the fraction of population altered within 100 generations compared to the equilibrium frequency, especially when alteration may be temporary (see Results). For organisms such as mosquitoes in the tropics, 100 generations would correspond to approximately a period of 10 years.

### Population suppression

Our model tracks the genotypic frequencies and fitnesses in each generation, thus allowing us to calculate the genetic load in each generation. As mentioned above, accurately estimating the level of actual population suppression across generations requires explicit incorporation of many ecological details. For instance, the nature of density dependence would be a critical factor in determining the level of population suppression that can be achieved through a given amount of genetic load (see Alphey and Bonsall 2014). However, suppression with any gene drive does require reasonably high genetic loads. Although incorporation of population dynamics is beyond the scope of this study, genetic load can be used to compare the *capacity* of each gene drive for population suppression (Burt 2003). As population fitness often is highly dependent on female fitness, payload genes that reduce only female fitness can be propagated through males and still achieve high genetic loads. We compare the genetic load generated by the three gene drives with payload genes that have a multiplicative or recessive female-limited fitness cost.

As migration adds another complex factor that may influence population suppression, we restrict our comparison of the capacity for population suppression only to the scenario of an isolated target population. Moreover, because a frequency-only model is not well suited for addressing the long-term effects of high genetic load on a population, we compare the mean genetic load achieved with a gene drive only within the first 20 generations. As described above, the genetic load in a generation is given by the ratio of mean absolute fitness in the population to the absolute fitness of a population with only wildtype homozygotes.

## Results

### Performance in isolated populations

We first compare the threshold introduction frequency needed for each of the gene drives to successfully spread a payload allele with a given fitness cost into an isolated population. The threshold introduction frequency is a useful measure of the initial effort required to conduct population alteration with a given gene drive. Moreover, it may also influence the ability of a gene drive to spread into new populations through migration (see localization results below). All three gene drives studied here exhibit a threshold for the introduction frequency, below which the drive fails to successfully spread the payload allele into a population. The threshold introduction frequencies are the lowest for the daisy-chain drive, highest for the one-locus underdominance drive, with the two-locus underdominance drive having relatively intermediate threshold introduction frequencies (Figure 3 A, B & C). For instance, a successful spread of a payload allele with a homozygous fitness cost of 10% requires a threshold introduction frequency of 3% (transgenic:wild-type release ratio ≈0.031:1) with the daisy-chain drive, while the two-locus and one-locus underdominance drives require introduction frequencies greater than 34% (transgenic:wild-type release ratio ≈0.52:1) and 72% (transgenic:wild-type release ratio ≈2.57:1), respectively. The reason for this difference is that the underdominance drives work through large reductions in fitness of individuals that carry only one of the gene drive constructs. This leads to a loss of a large fraction of gene drive alleles, especially when the gene drive is present at low frequency in the population. The threshold introduction frequency changes with the cost of the payload allele for all types of gene drives, with more costly payload alleles requiring progressively larger initial introductions (Figure 3 A, B & C).

**Figure 3:**
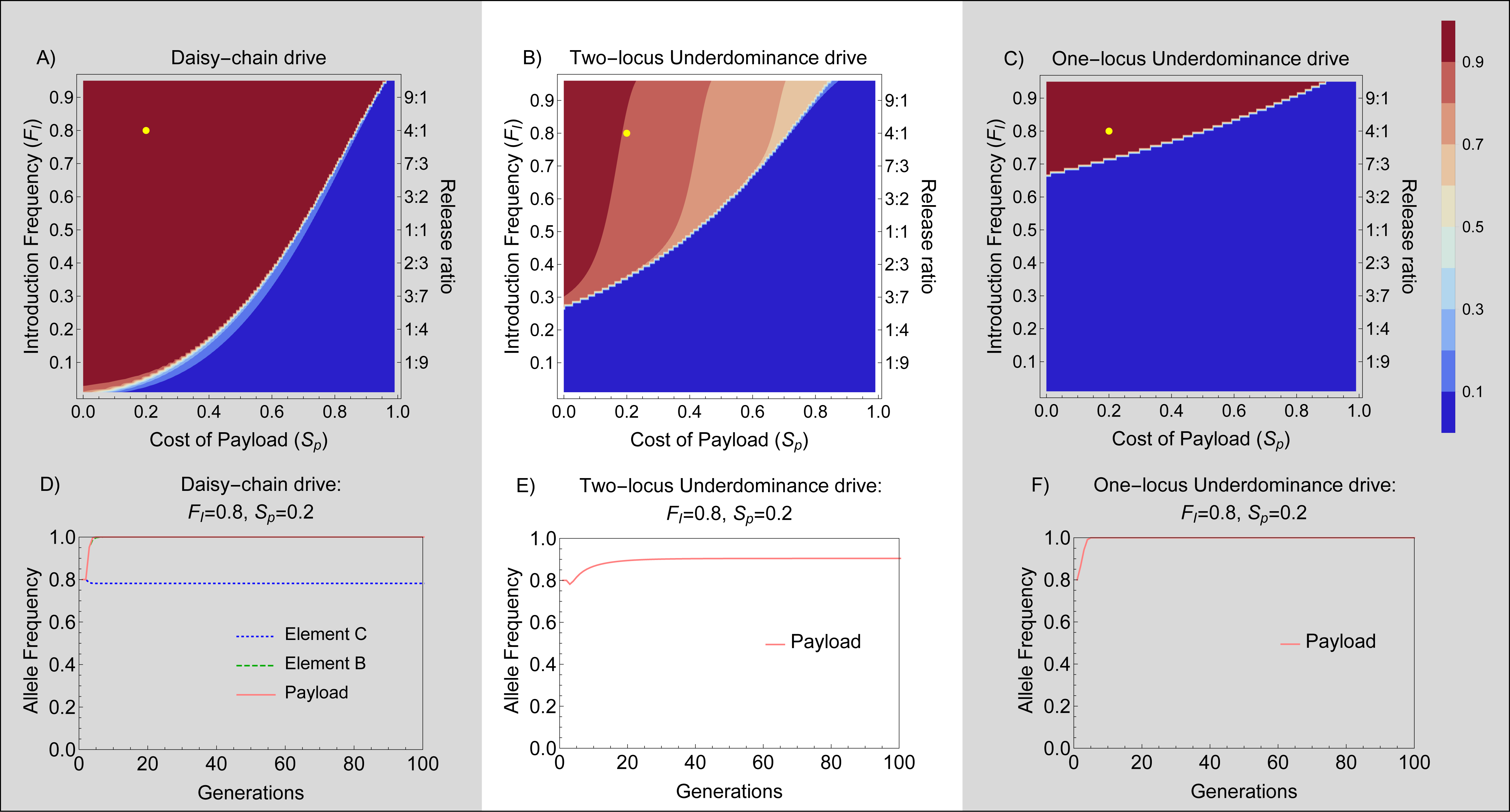
Performance in isolated populations – Contour plots show the mean frequency of the payload allele in the 100 generations following a single introduction of engineered organisms, with color values given by the bar legend. The vertical axis shows the introduction frequency (F*_I_*) on the left side and the corresponding release ratio on the right side of each panel. Horizontal axis shows the homozygous cost of the payload allele. A) Daisy-chain drive, homing efficiency=0.95; B) Two-locus underdominance & C) One-locus Underdominance. The yellow dot on each contour plot shows the conditions with which the time series plots (D, E & F) were generated.

Another important difference between the gene drives is that the daisy-chain drive and the one-locus underdominance drive do not have stable internal equilibria for the payload allele frequency. Both the drives either push the payload to near fixation (payload allele is maintained at a frequency within 10^−15^ of 1) or are eliminated from the population. The daisy-chain drive can, under a narrow set of conditions (contours between dark red and dark blue in Figure 3A), drive the payload to high frequencies with an eventual decline after the other two elements of the daisy-chain are lost. The two-locus underdominance drive, on the other hand, has an equilibrium payload allele frequency lower than 100% (Figure 3 B and E). Depending upon the cost of the payload allele, this equilibrium allele frequency may be as low as 70% (Figure 3 B, light pink areas). The equilibrium *genotypic* frequency of the payload allele at birth, i.e., the frequency of individuals carrying at least one copy of the payload allele at birth, however, does reach greater than 90% even when equilibrium payload allele frequency is close to 70% (i.e. the frequency of wildtype homozygotes is less than 10%). This is because most wild-type alleles at the A locus (the locus that can carry the payload allele) are present in a heterozygous state. Moreover, if the toxin produced by the underdominance drive is lethal during early (pre-adult) life stages, the equilibrium genotypic frequency of the payload (frequency of individuals carrying at least one copy of the payload) in the adult population practically reaches fixation (> 99.99%). This is because any individuals born homozygous for the wild-type allele at the A locus would die before reaching adulthood if they also possess a copy of the transgene at the B locus (note that the census for all allelic frequencies in our model occurs at birth). Depending on when lethality occurs, this difference between the drives can be critical for certain applications, where the maintenance of a few wild-type alleles in the population may be detrimental.

### Effect of unidirectional migration into the target population

The performance of all three types of gene drives in the target population decreases when wild-type individuals can migrate into the population (Figure 4). Migration generally increases the minimum number of organisms that need to be released for spreading a given payload gene into a target population. In the case of the daisy-chain drive, migration of wild-type individuals into the target population can also cause the spread of the payload gene to be temporary, with the population eventually reverting to a wild-type state (Figure 4 A, right side panels). Both underdominance drives, once established, are much more resistant to migration than a daisy-chain drive. When temporary alteration of a population is desired, this property of the daisy-chain drive may prove useful. It also suggests that in isolated populations, the daisy-chain drive would be more amenable to reversal, compared to the underdominance drives, allowing the population to be restored to a wild-type state if so desired. It should be noted, however, that the scenario shown in Figure 4 represents unidirectional migration into the target population, so that a source of wild-type individuals is maintained on the mainland. The gene drives differ in their level of localization (see below), and reversal in the target population would not be as feasible if surrounding populations had high frequency of the gene drive.

**Figure 4:**
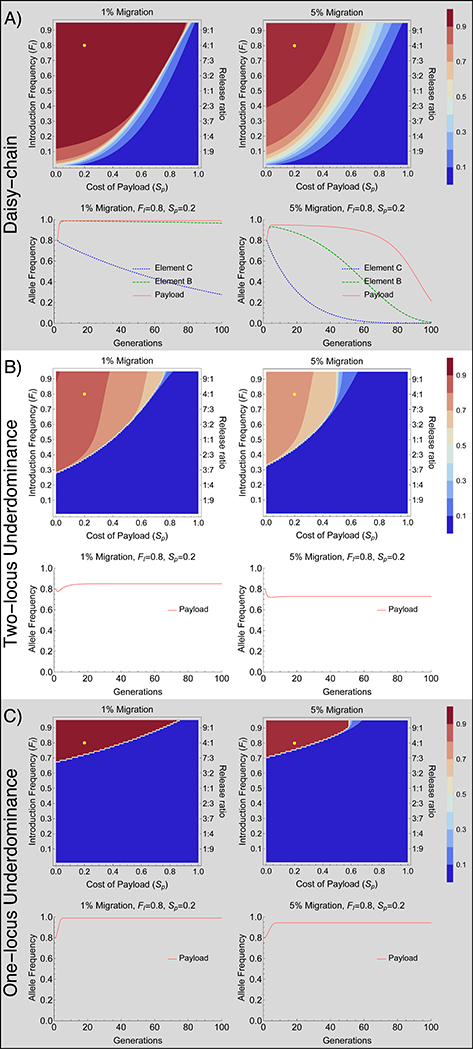
Effect of unidirectional migration – Contour plots show mean frequency of the payload allele in the first 100 generations after a single introduction of engineered organisms with migration only into the target population. The time series plots correspond to the conditions given by the yellow dots on the contour plots for the respective gene drives. Background shading highlights panels for different gene drives.

### Level of localization

To address the level of localization we follow the frequency of the payload allele in the target population and a neighboring population, with bidirectional migration. Genetically modified individuals are released only in the target population. The three types of gene drives differ considerably in their level of localization. Being the gene drive that requires the smallest introductions, the daisy-chain drive is also the least localized of the three gene drives, even with low levels of migration (Figure 5 A, D & G). That is, under most conditions when the daisy-chain drive successfully spreads the payload allele in the target population, it also spreads to high frequency in the neighboring population. The two-locus underdominance drive can achieve localized population alteration, when migration rates are low, but also requires relatively higher initial introductions of modified organisms (Figure 5 B, E & H). The one-locus underdominance drive requires the largest introductions and very low migration rates to spread successfully. But, when the one-locus underdominance does spread successfully in the target population, it remains highly localized (Figure 5 C, F & I) and never spreads to the neighboring population under any conditions that we studied.

**Figure 5:**
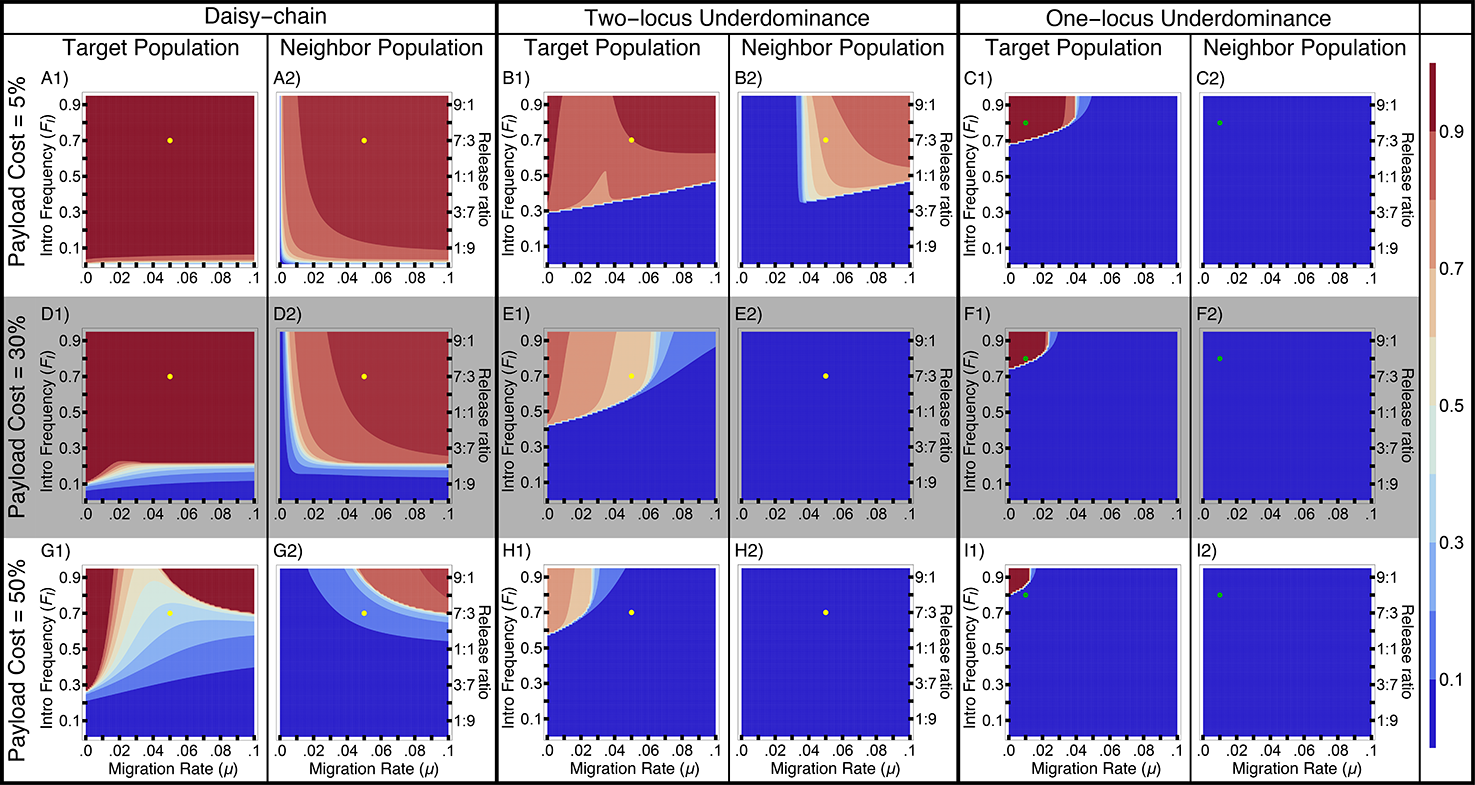
Localization analysis with bidirectional migration – Contour plots show mean frequency of the payload allele in the first 100 generations in the two populations, after a single introduction of transgenics into the target population. The dots on the contour plots are shown to facilitate visual comparison between figures for different payload costs (Yellow dots for daisy-chain and two-locus underdominance correspond to F*_I_* = 0.7 and *μ* = 0.05; green dots for one-locus underdominance corresponds to F*_I_* = 0.8 and *μ* = 0.02). Payload costs given for each row are the costs incurred by individuals homozygous for the payload transgene.

The level of localization changes with the fitness cost of the payload gene. In an isolated population, a gene drive carrying a less costly payload gene can spread more easily than a drive carrying a costly payload because of the higher threshold introduction frequencies of costly payloads (see Figure 3). But this also causes gene drives with low-cost payloads to be less localized than gene drives with costly payloads when migration occurs between two populations (Figure 5). Higher threshold introduction frequencies make it harder for the gene drives to establish in the neighboring population. Another reason for the pronounced pattern of higher localization with increasing payload cost is that high payload costs impose higher genetic load on the target population, which in our model reduces the effective immigration rate into the neighboring population from the target population, and increases the effective immigration rate of wild-type individuals coming into the target population. This pattern still holds, and becomes only slightly less pronounced, when migration rates are independent of genetic load (Appendix 3).

As mentioned in the Model section above, we assume that migration is independent of individual genotype. In reality, high payload costs may result in genotype-dependent reduction in migration rates of individuals carrying the gene drive. In general, localization is likely to be harder to maintain for gene drives designed for population replacement, which are likely to have low payload costs.

Migration, in general, tends to restrict the spread of the gene drive in the target population by constantly adding wild-type individuals. However, very high rates of bidirectional migration can result in the gene drive becoming established in both populations, removing the source of wild-type individuals. This is apparent in that intermediate rates of bidirectional migration are more restrictive for the establishment of the gene drive in the target population than very high rates under certain conditions for the daisy-chain and the two-locus underdominance drives (see Figure 5 B1 & G1). It is important to note here that we address the spread of a payload gene only to a population directly exchanging migrants with the target population. The spread to a secondarily connected population, once removed from the target population (i.e. a population that exchanges migrants with the neighbor population but not with the target population) is likely to be much smaller. This may allow a source of wild-type individuals to be maintained.

### Population suppression

We compare the mean genetic load that can be imposed upon isolated populations with the three gene drives. The population genetic load depends upon the fitness cost of the gene drive as well as the frequency of the gene drive alleles. Gene drives with a multiplicative female-limited payload gene expression yield higher genetic loads than those based on a recessive payload expression (see Appendix 4). The daisy-chain drive allows high mean load to be achieved with smaller introductions than the underdominance drives (Figure 6). For species that exhibit negative density-dependent dynamics, genetic loads close to or above 80% are desirable for significant population suppression (Burt 2003). Both underdominance drives addressed here allow such high loads only with very large introductions (Figure 6).

**Figure 6:**
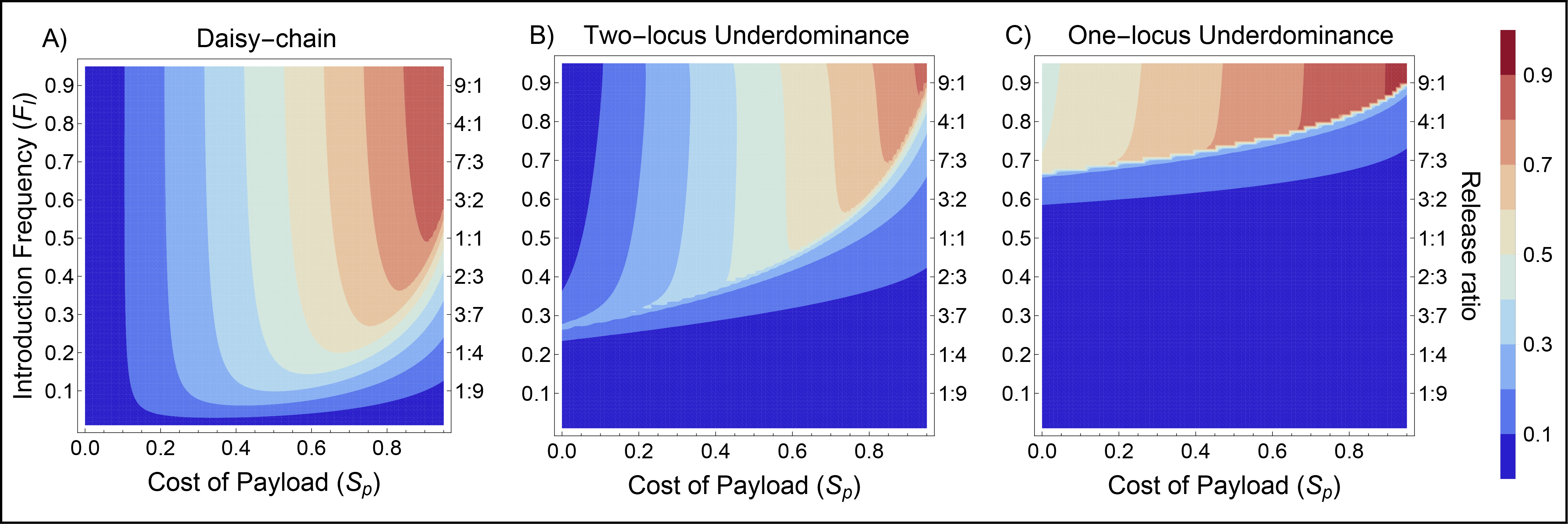
Mean load in isolated populations – Contour plots show mean of genetic load on an isolated population within the first 20 generations after a single introduction of gene drives with multiplicative female-limited payload cost.

## Discussion

There have been previous theoretical explorations of the dynamics of underdominance-based gene drives and of the daisy-chain drive (Davis et al. 2001; Magori and Gould 2006; Marshall and Hay 2012; Noble et al. 2016). Because these studies were done separately, and were not intended to make comparisons between different drives, each gene drive was analyzed under a different set of conditions. Our goal in this study was to compare the level of localization and the permanence of these gene drives under identical scenarios. Our modeling is limited to one or two populations, and does not include an exploration of ecological details such as different potential forms of density dependence. However, it enables some general predictions about the comparative characteristics of the three gene drives analyzed.

Overall, in isolated populations, the daisy-chain system is theoretically capable of driving a payload (the desired gene or gene disabling construct) to fixation with a release of far fewer engineered individuals than are needed for either of the underdominance drives. Compared to the one-locus system, the two-locus underdominance drive requires a lower threshold introduction frequency for drive to occur. In addition to the differences in the threshold introduction frequencies required for each gene drive, they also differ in the extent to which they alter an isolated population even when successful drive does occur, which may affect their utility for different applications. The daisy-chain system does not have a stable internal equilibrium in the scenarios studied here (i.e. payload either goes to fixation or is completely removed from the population). However, within a very narrow range of fitness costs and release numbers (see Figure 3A – region between dark red and dark blue contours), the daisy-chain system could temporarily drive a payload to a high frequency (but not fixation) with eventual loss from the population. Under this narrow set of conditions, the daisy-chain system results in an incomplete and temporary population alteration. When there is a fitness cost to the payload, the two-locus underdominance system is not expected to reach fixation even when drive occurs (i.e. it has a stable internal equilibrium; but see the discussion of dominant payload costs in Appendix 1). This property may affect the suitability of the two-locus underdominance system for certain applications, if the payload is not effective in heterozygous condition, or if the maintenance of a low frequency of wild-type homozygous individuals can pose a problem. The one-locus system reaches fixation as long as engineered individuals are released above the threshold introduction frequency for drive to occur.

Once the single-locus underdominant drive system reaches fixation, eliminating it from an isolated population would require release of a large number of wildtype individuals, or release of engineered individuals that disable the drive system (e.g. Individuals with an independently assorting transgene that suppresses the promotor of the toxin gene and itself has no or low fitness cost). Eliminating a two-locus underdominance drive after it reaches an equilibrium above the threshold introduction frequency would require similar releases. Additionally, over time the intermediate equilibrium frequency could enable recombination events that disable the two-locus underdominance drive and/or result in breakdown of the payload (Beaghton et al. 2017). The number of wild-type individuals needed to reverse the daisy-chain system depends upon the frequencies of the non-payload elements of the drive, which in turn depend upon their costs, the initial release size and time since the initial release.

The situation is more complex when there is one population, targeted for gene drive, and a second non-target population which exchanges migrants with the target population. In certain scenarios, migration may only be unidirectional into the target population; for example, if the target population is a small island near a much larger, randomly mating continental population, or if an aquatic target population is isolated from a non-target population by a barrier, such as a dam or a large waterfall, which restricts upstream migration. In such a case any migration to the target population elevates the number of engineered individuals that need to be released for drive to occur, and for the two-locus underdominance system, it also greatly decreases the equilibrium frequency of the payload. For the daisy-chain drive, the ongoing migration insures that the payload will eventually be lost from the island population, even if payload frequencies reach and remain at very high values for many generations.

In many real scenarios migration is likely to be bi-directional. This would be the case with two islands of similar size or with two mainland populations isolated by distance and/or physical barriers. If base migration rates are similar in both directions, the daisy-chain drive is predicted to become fixed in both populations under most conditions, unless payload costs are quite high. For the two-locus underdominance system, spread to the non-target population does not occur at 30% payload fitness cost, but can occur at specific release and migration rates when the payload costs are low (close to 5%). The one-locus underdominance system is the most localized of the three systems, and does not spread to the non-target population, even when the payload cost is 5%. Thus concern about unintended spread is highest with the daisy-chain system and lowest with the one-locus system. It must be noted that we did not examine spread to a secondarily connected population, one that does not exchange migrants with the target population, but does so with the ‘neighbor’ population. One possibility is that as the daisy-chain system spreads into secondarily connected populations, each subsequently connected population may receive fewer elements of the daisy-chain, or receive them at progressively smaller frequencies relative to the payload. However, the equilibrium frequencies for each given element of the daisy-chain in the two populations that are modeled here are equal, giving no indication of sequential removal of daisy-chain elements across populations.

In our models we always assume only a single release of the engineered individuals. Depending upon the specific application, multiple smaller releases, as examined by Davis et al. (2001) and Magori and Gould (2006), could impact the comparison. However, among the gene drives addressed here, it seems to be an inescapable fact that gene drives that require lower threshold introduction frequencies for drive to occur are also going to have higher risk of unintended spread through migration.

One factor that may greatly influence the localization of gene drives to target populations is genotype-dependent variation in migration rates. In our analyses we assume that the migrating group of individuals is perfectly representative of the source population. If the payload or other components of a gene drive reduce the likelihood of migration for the individuals bearing them, the risk of unintended spread would be greatly reduced compared to the results shown here. Genes that reduce the mobility of organisms, e.g. through physical deformities or by reducing desiccation tolerance, may therefore prove ideal for designing gene drives that can reduce population fitness and also have a lower risk of unintended spread.

Our analysis of localization in this study was limited to a two-population scenario, with no heterogeneity between the populations. The dynamics of these gene drive systems would certainly be different in a scenario with a more complex spatial structure with multiple populations spread across two dimensions. For one, the target population may receive more wild-type immigrants, which may make all three gene drives more difficult to establish in the target population. The exact dynamics would depend upon the particular two-dimensional pattern of migration between the different populations (e.g. Láruson and Reed 2016). However, inclusion of more populations by itself may not change our results qualitatively – gene drives with lower threshold introduction frequencies would still be more likely to spread to neighboring populations. Studies with more complex spatial structure would still be very useful in understanding the behavior of these gene drives in natural populations, and would be a fertile avenue for future studies.

In isolated populations, all three drive systems are able to spread a payload into a population as long as the fitness cost associated with it is low, as could be the case if the payload was built to change a characteristic such as ability to transmit a pathogen. However, if the goal of a release is to suppress an isolated population to very low density or cause local elimination, the daisy-chain drive with a female-limited payload cost can be much more efficient than the underdominance drives. Migration in one or two directions is likely to reduce the capacity of the gene drives for suppression, because migration reduces the ability of a gene drive to spread in the target population, and even after successful suppression, migration may restore the population.

The general models presented here could be expanded to include more population ecology and structure, and parameters could then be adjusted to fit specific targeted populations. It would also be useful to expand modeling efforts to include economic estimates regarding the financial and time costs involved in development of each of these drive systems. Given the genetic engineering tools available today, the cost for developing a daisy-chain system is likely to be greater than the costs for developing the underdominance systems. For example, the daisy chain system requires that at least two chromosomes be engineered to express multiple guide RNAs, and that the targets of the guide RNAs on at least two chromosomes are essential genes that are haplo-insufficient. Moreover, it must be possible to insert a construct with the replacement haplo-insufficient gene along with Cas9 and guide RNAs without a substantial fitness cost (Noble et al. 2016). For a pest species without highly developed genomic tools, these requirements could be hard to achieve without very high development costs. We model the impacts of a 10% fitness cost of these non-payload insertions (Figure S7), but if even the cost of one of the elements in the wild turns out to be high (50%), the system efficiency declines (Figure S8 and S9). Both underdominance systems are also complicated to construct, and can have fitness costs due to leaky toxin gene expression (Akbari et al. 2013). But, because fewer insertions are involved and expression can be in any tissue/time, they currently appear to face fewer engineering hurdles.

Another factor that our analysis does not consider is the evolution of resistance to the gene drives. Wild-type alleles resistant to CRISPR-Cas9-based homing drives can be present in genetically diverse populations, and can arise through non-homologous end-joining (Reed 2017; Drury et al. 2017; Champer et al. 2017a; Champer et al. 2017b). Although the daisy-chain drive design includes multiple guide-RNAs and the requirement for haplo-insufficiency of the genes, it is not yet clear how effective these measures will be for preventing resistance to the drive in natural populations. Recent studies of homing drives with multiple guide-RNAs suggest that that this design does reduce, though not eliminate, the formation of resistant alleles (Noble et al. 2017; Champer et al. 2017a). Evolution of resistance to the two-locus underdominance drive is also possible to evolve, though it is perhaps more difficult than evolution of resistance to CRISPR-based gene drives. For example, evolution of suppressors that are not linked to any toxin can disable an underdominance drive. Evolution of resistance will certainly reduce drive efficiency, and depending upon the distribution of resistance alleles in different populations, it may even affect the level of gene drive localization. Presence of resistant alleles in the neighboring population may improve gene drive localization, or may result in drive failure if migration brings sufficient resistant alleles into the target population. Indeed, researchers are continually trying to design gene drives that can overcome the issue of resistance (e.g. Esvelt et al. 2014; Noble et al. 2017; Champer et al. 2017a). Our goal in this study was to compare the designs of these gene drives assuming the drives function according to the design. As more empirical data becomes available about the performance of these gene drives against resistance, and the prevalence of resistance in natural populations, more detailed models that include the evolution of resistant alleles would be needed.

A number of authors and committees have emphasized the need for development of spatially limited gene drive systems (e.g. Gould et al. 2008; Marshall and Hay 2012, NASEM 2016). The systems described here could meet that call under certain conditions if the goal of the gene drive is to change population characteristics. A thorough investigation of the ability of these drives to carry out localized population suppression will need further modeling that incorporates species specific population dynamics. Which of these gene drives would be the best choice for a specific target population would depend in part on the balance between concern over unintended spread, need for local permanence, cost of producing the drive and cost of rearing the necessary number of released individuals.

## Acknowledgements

We are grateful to B. Hollingsworth, J. Baltzegar, J. Kang, J. Sudweeks, C. Gunning, K. M. Esvelt, M. Edgington, the associate editor and three anonymous reviewers for comments on earlier versions of this manuscript. SD was supported by a grant from the W. M. Keck Foundation to FG. MRV was supported by IGERT fellowship through the National Science Foundation. FG and ALL were funded by National Institute of Health grant R01-AI091980 and a National Science foundation grant RTG/DMS – 1246991.

## Data archiving statement

Mathematica code for the models will be made available by request upon acceptance of the manuscript for publication.

